# Magnetic Microtweezers for High-Throughput Bioseparation in Sub-Nanoliter Droplets

**DOI:** 10.1101/2024.01.18.576187

**Authors:** Simon Dumas, Lucile Alexandre, Mathilde Richerd, Marco Serra, Stéphanie Descroix

## Abstract

Multiomics studies at single-cell level require small volume manipulation, high throughput analysis and multiplexed detection, characteristics that droplet microfluidics can tackle. However, the initial step of molecules bioseparation remains challenging. Here, we describe a unique magnetic device to trap and extract magnetic particles in sub-nanoliter droplets, for compartmentalisation of detection steps. Relying on electrodeposition of NiFe structures and microfluidic manipulation, this technology has allowed the purification of genetic material at single-cell scale and was able to reach an extraction rate of 72% for a sample of purified oligonucleotides.

## 1. Introduction

Miniaturization allows playing with the kinetics of reactions, reducing time of analysis, volumes of sample and reagents, risks of contamination coupled with increasing the sensitivity of detection. Development of miniaturized systems has supported this shift, reinforced by a better comprehension of the physics at micro-scale [1]. Microfluidics often finds applications in diagnostics as it offers the possibility to work with small quantities of reagents and samples at limited costs [2]. The fabrication of micro-tools enables the implementation of various steps of laboratory work in microfluidics systems, and translation of fastidious protocols to a single instrument.

Working with a volume range from microliters to picoliters can be coupled with compartmentalization of reagent via droplet-based microfluidics [3], [4]. As many microfluidic approaches, droplet encapsulation allows the manipulation of small samples in a confined environment, reducing time of analysis and contamination risks, opening the door to scale-up, parallelization and multiplexing. Droplet microfluidics have progressed toward high impact studies with complex applications [5] at high throughput such as biomarkers detection [6], drug screening [7], 3D cell culture [8], [9] or single-cell measurements [10], [11]. Generation, merging, mixing, sorting and splitting of droplets can be achieved at microscale, relying on micro-fabrication and surface treatment development [12]. However, some operations remain challenging to achieve such as extraction and purification steps, as they often rely on a physical separation.

To tackle this challenge, the magnetic tweezers technology has been developed by our team for the extraction and redispersion of magnetic beads inside droplets [13]. The versatility of a magnetic solid-phase extraction makes it an obvious candidate for translation to micro-scale. The technology of magnetic tweezers relies on the control by tunable electric coil of an external magnetic field focused on the path of droplets. Magnetic particles inside the droplet are then confined in a compact cluster that can be extracted from the initial droplet, washed and released in a second droplet by switching off the magnetic field. This process has already shown its efficiency for biomarkers extraction and detection [14]–[16].

Here, we present a new generation of magnetic tweezers for the manipulation of magnetic micro-particles in sub-nanoliter droplets and its application to purification of small quantities of mRNA compatible with single cell analysis. This new approach is based on NiFe micro-tweezers, which are fabricated by photolithography and electrodeposition, and are actuated by an external permanent magnet. Modification of the geometry and structure of the micro-tweezers tackle the challenge to work at high throughput with sub-nanoliter droplets without sacrificing the sensitivity. This new design has allowed us to work at single-cell level, with 0.5 nL droplet at a throughput of 20 Hz [17]. This unique magnetic configuration has bridged the gap between high throughput microfluidic sample manipulation and multi omics analysis at single cell level.

## 2. Materials

### 2.1. Materials for substrate preparation before plating

1. Large microscope glass slides 50 x 75 mm (Corning)
2. Sputtering machine equipped with Cu and Ti targets
3. Photolithography equipment (chromium mask, mask aligner, hotplates, spin coater)
4. Photoresist adhesion promoter (TI-Prime, Microchemicals)
5. Negative photoresist (AZ 125 nXT, Microchemicals) and developer (AZ 726 MIF, Microchemicals)

### 2.2. Materials for electroplating

1. Prepare plating solution by dissolving in 700 mL of water the following reagents: 250 g NiSO_4_ · 6H_2_O, 5 g FeSO_4_ · 7H_2_O, 25 g Boric acid, 2 g Saccharin, 0.1 g Sodium dodecyl sulfate. Place the beaker on a hotplate at 40 °C while stirring until complete dissolution, and fill with water up to 1 L. Filter the solution through Stericup (Millipore). The solution must be dark green and can be kept and reused for at least 3 months at room temperature.
2. DC generator (ALR3002M, ELC)
3. Conductive copper tape (1181/9.5, 3M)
4. Pure nickel anode, 100 x 60 x 2 mm (Goodfellow)
5. UV Ozone cleaner (Jelight)
6. Arduino Uno
7. Relay board (Radiospares, 843-0834)
8. Photoresist stripping solution, Technistrip P1316 (Technic)

### 2.3. Materials for microfluidic chip preparation

1. Photolithography equipment (chromium mask, mask aligner, hotplates, spin coater, SU-8, developer)
2. PDMS and curing agent (Sylgard)
3. Mask aligner (MJB4, Süss) or custom tabletop aligner (Li et al., *Rev. Sci. Intrum*., 2015; Guglielmotti et al. *HardwareX*, 2022).
4. Tetraethyl orthosilicate (TEOS)
5. 1H,1H,2H,2H-Perfluorooctyl Trichlorosilane

### 2.4. Materials for microfluidic device operation

1. Syringe pumps (Cetoni)
2. Bright field inverted microscope
3. Biopsy punches (diameter 0.75, 6 and 8 mm)
4. 1 mL syringes (Omnifix, Braun) and beveled needles 23G and 27G (Terumo)
5. PTFE tubings (Adtech BIOBLOCK/05 and BIOBLOCK/13)
6. Neodymium magnets (5 mm cube, N50, and 10 mm cube, N42, Supermagnete)
7. Carrier oil (HFE-7500 with 2% fluorosurfactant, Emulseo)

### 2.5. Materials for mRNA extraction in droplets

1. Streptavidin Dynabeads MyOne C1 (Thermofisher, 65001)
2. Dynabeads solution A (0.1 M NaOH, 50 mM NaCl)
3. Dynabeads solution B (100 mM NaCl)
4. 2X Binding/Wash buffer (10 mM TrisHCl pH7.5, 1 mM EDTA, 2M NaCl, 0.05% *v/v* Triton X-100)
5. 1X Binding/Wash buffer (Dilute 2X buffer in same volume of water)
6. Biotinylated oligo-dT30VN (5’-/5BiotinTEG/AAGCAGTGGTATCAACGCAGAGTACTTTTTTTTTTTTTTTTTTTTTTTTTTTTTTVN)
7. Cell Resuspension Buffer (15% *v/v* Optiprep, 0.75% BSA in PBS)
8. 1.5X Lysis/Binding Buffer (150 mM TrisHCl pH7.5, 750 mM NaCl, 15 mM EDTA, 0.075% *w/v* Triton X-100, 0.75% *w/v* BSA, 1 U/µL Ribolock RNAse Inhibitor). Prepare fresh and keep on ice until use.
9. Emulsion breaking solution (20% *v/v* 1H,1H,2H,2H-perfluoro-1-octanol in HFE-7500)

### 2.6. Materials for mRNA quantification

1. qPCR equipment (Quantstudio 3, Applied Biosystems)
2. RT-qPCR kit (CellsDirect One-Step qRT-PCR, Thermofisher, 11753100)
3. ACTB qPCR primers (Fw 5’-GGATGCAGAAGGAGATCACTG, Rv 5’-CGATCCACACGGAGTACTTG)
4. ACTB TaqMan fluorescent probe (5’-/56-FAM/AAGATCAAG/ZEN/ATCATTGCTCC/3IABkFQ/)

## 3. Methods

The microfabrication of substrates and chips requires a clean room environment and special equipment to perform photolithography and physical vapor deposition (PVD). This includes a chromium mask, a mask aligner, a spin coater, hotplates, a sputtering machine, a plasma chamber, a vacuum desiccator. First, the magnetic micro patterns are fabricated using photolithography and electroplating on glass slides. Second, the microfluidic chip is obtained by PDMS molding on a SU-8 patterned wafer, and is aligned and bonded onto the magnetic patterns.

### Design considerations

The designs of the chip and the patterns must be complementary, and a chamber must be provided to accommodate the magnets. For the pattern design, an area for connecting the alligator clip during electroplating must be anticipated. This area must remain dry during electroplating, and is preferably placed at least 1 cm away from the patterns.

### 3.1. Preparation of the substrates for electroplating

1. Clean a glass slide with acetone, isopropanol, and proceed to seed layer deposition by sputtering of 20 nm Ti and 200 nm Cu (**Note 1**).
2. Apply TI-Prime by spin coating at 4000 rpm for 30 s, and dry the substrate on hotplate at 95°C for 1 min.
3. Apply negative photoresist AZ 125 nXT by spin coating (**Note 2**), and bake for 4 min at 95°C, 10 min at 125°C. Let cool down to room temperature.
4. Using a mask aligner, expose the substrate to 2 J/cm^2^ of UV-light (i-line, 365 nm) through a chromium mask. Develop the photoresist by immersion in AZ 726 MIF for 2 min, rinse with water and dry. The substrate can be kept protected from light for at least three months.

### 3.2. Electroplating of NiFe structures

This step may require optimization and various problems can be encountered. Several parameters can significantly influence the result in terms of deposit quality, magnetic properties or thickness. They include deposition time, electrical current, patterns design, stirring speed, bath temperature and pH, presence of dust in the bath, evaporation. The protocol was adapted from [18] and modified for higher thicknesses. It was observed that the deposit will detach after long periods of deposition. For this reason, a programmable switch was added to periodically open and close the circuit, decreasing the internal stress within the deposit.

1. Place a beaker containing around 700 mL of plating solution on a hotplate set at 40 °C with continuous stirring.
2. Perform UV-Ozone for 10 minutes on the substrate to improve patterns wettability, and wet them using a wash bottle\ filled by 1% SDS.
3. Apply copper tape on the substrate (using a dedicated dry area) and on the nickel anode, and connect them with alligator clips.
4. Connect them to the DC generator, the substrate to minus and anode to plus, with the relay used as an interrupter to programmatically open and close the circuit. Connect the relay to the Arduino: GND pin to GND, power pin to +5V, and the control pin to digital output 2.
5. Immerse the substrate and the anode in the bath, they must face each other with a distance of 5-7 cm, using a 3D printer holder. Immediately turn the generator on at minimum power. Increase the current to reach a current density of 7 mA/cm^2^ (**Note 3, 4**).
6. Fill the beaker with a plating solution until all the patterns are immersed. Ensure that the copper tape and the alligator clip remain dry.
7. Power the Arduino board up to launch the following program loop:

~~~
int digout = 2; // Use digital output 2 on Arduino board
int counter = 0;
void setup() {
  pinMode(digout, OUTPUT);
}
void loop() { // Every 5 min, make a 15 s break. Every hour, make a 2 min break.
  counter = 0;
 while(counter != 12) {
  compteur ++;
  digitalWrite(digout, LOW);
  delay(300000); // 5 min
  digitalWrite(digout, HIGH);
  delay(15000); // 15 s
 }
 delay(105000); // 2 min
}
~~~
8. Let the substrate in the plating solution with continuous stirring (**Note 5**) until the thickness reaches 35 µm. The expected deposition time is 12 h, with a rate of 3 µm/h (**Note 6, 7**).
9. Clean the substrate with water, and immerse in TechniStrip P1316 at 70 °C while stirring until the photoresist and copper dissolve completely. This step can be repeated after 20 min by renewing the solvent to remove eventual residues (**Note 8**).

### 3.3. Preparation of the microfluidic chips

The fabrication of the PDMS counterpart is performed by conventional soft lithography, using 4” wafers and SU-8 2035 to achieve a thickness of 35 µm. PDMS with a curing agent ratio of 1:10 is used. This method is not detailed here as it is extensively described in the microfluidics literature.

1. Prepare the PDMS chips by soft lithography. To facilitate a precise alignment with the magnetic patterns, the PDMS slabs must be as flat as possible. This can be achieved either by leveling the oven during PDMS curing, or by curing the PDMS between two parallel plastic plates [19].
2. A thin titanium layer remains on the magnetic substrate, which inhibits plasma bonding with the PDMS, and the magnetic substrate needs to be treated with a silane. Activate the magnetic substrate with O_2_ plasma, and place it under vacuum in the presence of tetraethyl orthosilicate (TEOS) for 30 min.
3. Activate PDMS and magnetic substrate with O_2_ plasma before alignment.
4. Align and bond the two parts together using either a mask aligner [19] or a custom tabletop aligner [20], [21].
5. Press gently on the PDMS and place the bonded chip in an oven (or hotplate) at 70 °C for at least 10 min to ensure complete bonding.
6. The channels must be silanized to render them hydrophobic: prepare and filter a solution of 1% *v/v* 1H,1H,2H,2H-Perfluorooctyl trichlorosilane in HFE-7500 and flow it in the microfluidic channels using a syringe. After no more than 5 min, rinse with pure HFE-7500, blow air, and place back the chips at 70 °C to evaporate remaining oil. Chips can be stored for months at room temperature, protected from dust.

### 3.4. Preparation of collection/reinjection reservoirs

1. Using a 8 mm biopsy punch, cut a cylinder from a 5-10 mm thick PDMS slab.
2. Punch two holes using a 0.75 mm biopsy punch, clean with isopropanol.
3. Using a tweezer, pass a 0.56 mm (I.D.) tubing through one hole and a 0.3 mm (I.D.) tubing through the other.
4. Using a tweezer, push the PDMS plug in a 1.5 mL microcentrifuge tube.
5. Using a tweezer, push or pull the tubings so that the wider one reaches the bottom of the tube, and the thinner one end is at the PDMS interface (**Figure 2B**).
6. Prepare a syringe filled with carrier oil with a 23G needle, connect it to the wider tubing using a tweezer. Push the plunger to fill the reservoir until the oil drops from the thinner tubing. Keep the syringe connected to the reservoir.

**Figure 1.**
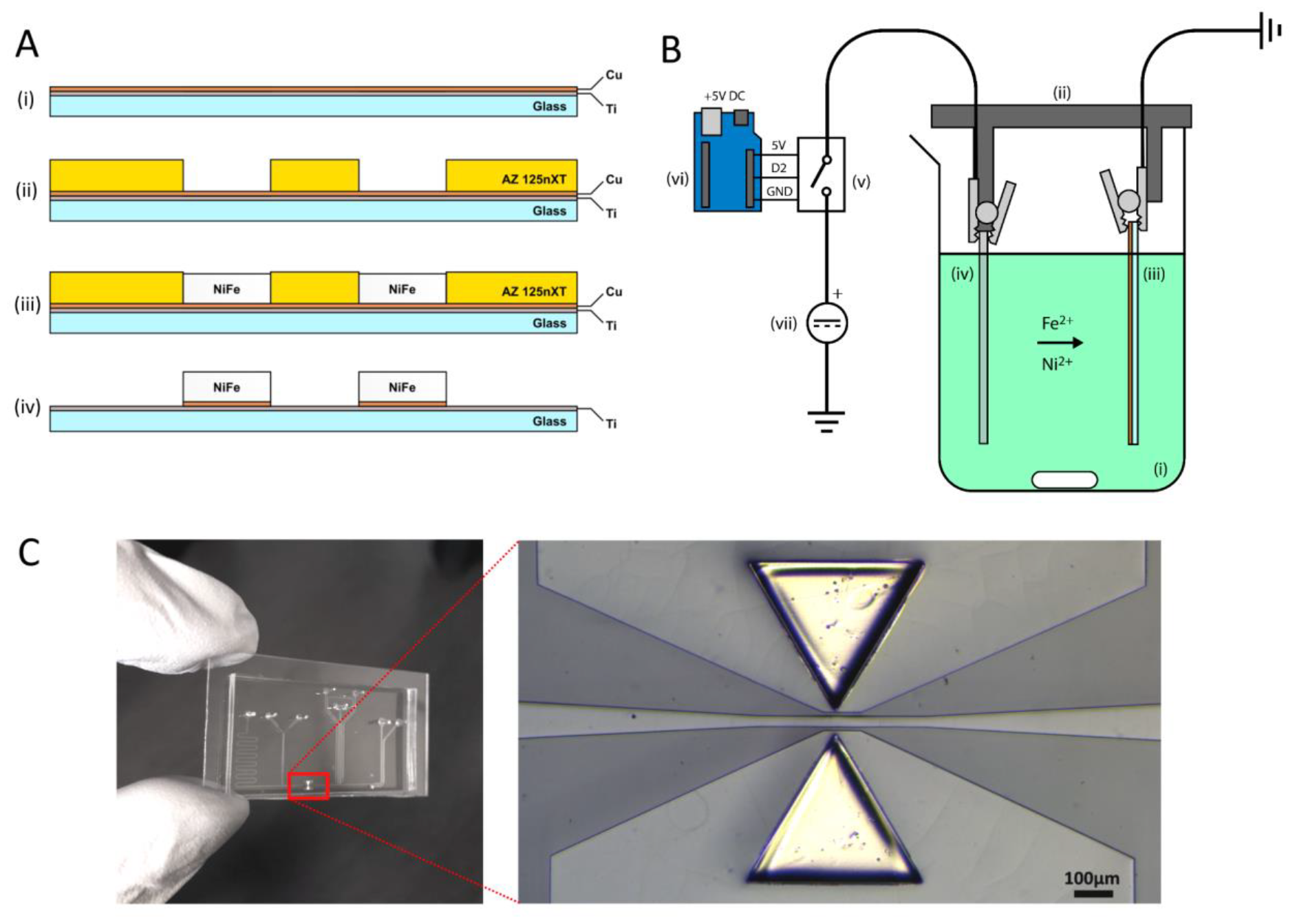
(a) Fabrication of the micromagnet substrate (i) Sputtering of a Ti:Cu layer on glass slide (ii) Photolithography (iii) Electroplating (iv) Stripping of photoresist and Cu, and TEOS treatment. (b) Electroplating process (i) Beaker filled with the plating solution (ii) 3D printed holder (iii) Substrate (iv) Nickel anode (v) Relay used as a programmable switch (vi) Arduino board (vii) Stabilized DC source (c) Final chip after alignment and bonding of the PDMS channels on the micromagnet substrate.

**Figure 2.**
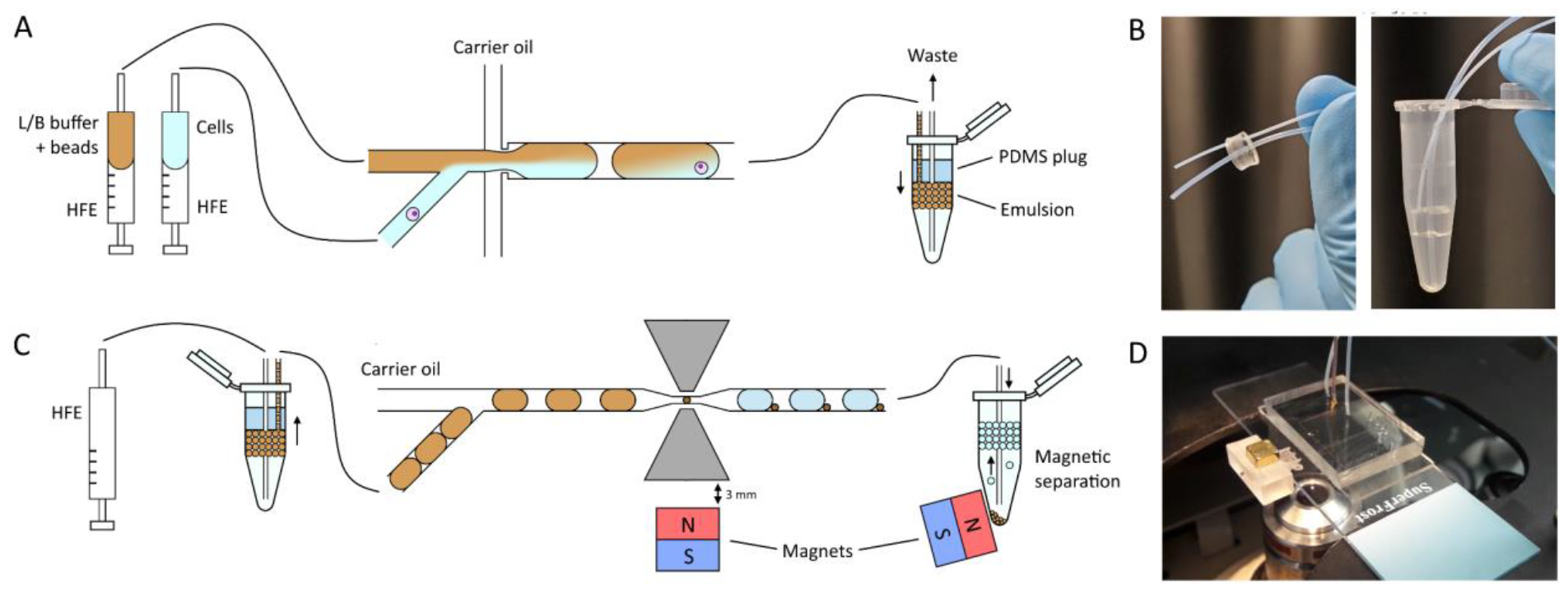
Microfluidic chip operation (a) Single cell encapsulation with beads and lysis buffer (b) Preparation of the collection/reinjection reservoir. (c) Reinjection of the droplets in the magnetic microtweezer chip. (d) Permanent magnet positioning using a custom 3D printed holder.

### 3.5. Preparation of oligo-dT conjugated beads

The extraction protocol is optimized for Dynabeads MyOne (Thermofisher). These beads are not commercialized with oligo-dT functionalization, and need to be conjugated in another way.

1. Take 80 µL of Dynabeads MyOne Streptavidin C1 (80 µg of beads)
2. Using a magnet, wash two times in 1 mL of Dynabeads solution A
3. Perform two additional washing with Dynabeads solution B
4. Discard supernatant and add 80 µL of 2X Binding/Wash buffer
5. Add 80 µL of 100 µM biotinylated oligo-dT30VN
6. Incubate 20 min at room temperature with rotation
7. Wash three times with 1X Binding/Wash buffer

### 3.6. Droplet generation and cell encapsulation

For cell encapsulation, a drop maker with two aqueous inlets is required. In this part, droplets of 0.5-1 nL are generated and stored in the reservoir. The droplets contain: cells, a lysis/binding buffer and oligo-dT magnetic beads (**Figure 2A**).

1. Harvest cells using a conventional cell culture protocol (not detailed here).
2. Resuspend the cells in 70 µL of Cell Resuspension Buffer at a concentration of 600 cells/µL.
3. Wash the oligo-dT conjugated beads with 1.5X Lysis/Binding Buffer and resuspend them in 200 µL of the same buffer at a concentration of 60 µg/µL. (**Note 9**)
4. Fill two syringes half way with HFE-7500 and with the above mentioned solutions. Mount the two syringes on the syringe pump vertically to allow the aqueous phase to remain above (**Note 10**). Mount a third syringe filled with carrier oil on the syringe pump.
5. Place the encapsulation microfluidic chip under a microscope. Plug 27G needles on the syringes and connect them to the chip with 0.3 mm (I.D.) tubings.
6. Generate 0.5 nL droplets by using the following flow rates: 200 µL/h for the buffer with beads, 100 µL/h for the cells, and 200 µL/h for the carrier oil. (**Note 11**)
7. When the generation is stable, connect the collection/reinjection reservoir to the chip, and disconnect the syringe at the reservoir outlet. The emulsion should fill the reservoir from the top. Keep the reservoir on ice or in a cold block during the whole experiment.
8. When the generation is finished, disconnect the reservoir from the chip and connect its outlet back to the syringe. Keep the reservoir on ice.

### 3.7. Magnetic beads extraction through the magnetic microtweezers

In this part, the emulsion is reinjected into the chip with the micro tweezers. The droplets are spaced with oil and the beads are extracted by the tweezers. After the extraction, the droplets and the extracted bead clusters are collected and separated in an external tube equipped with a permanent magnet. (**Figure 2C**)

1. Place the extraction microfluidic chip under a microscope with the 5 mm cubic magnet facing the tweezers at a distance of 3 mm using a 3D printed holder (**Figure 2D**).
2. Mount the reservoir syringe to the syringe pump, and connect the reservoir to the chip.
3. Inject the emulsion and spacing oil at 40 µL/h and 20 µL/h respectively. The droplet reinjection frequency should be around 40 droplets/s. The extraction of magnetic beads takes place at the micro tweezers. **Figure 3** shows what a good extraction looks like.

**Figure 3.**
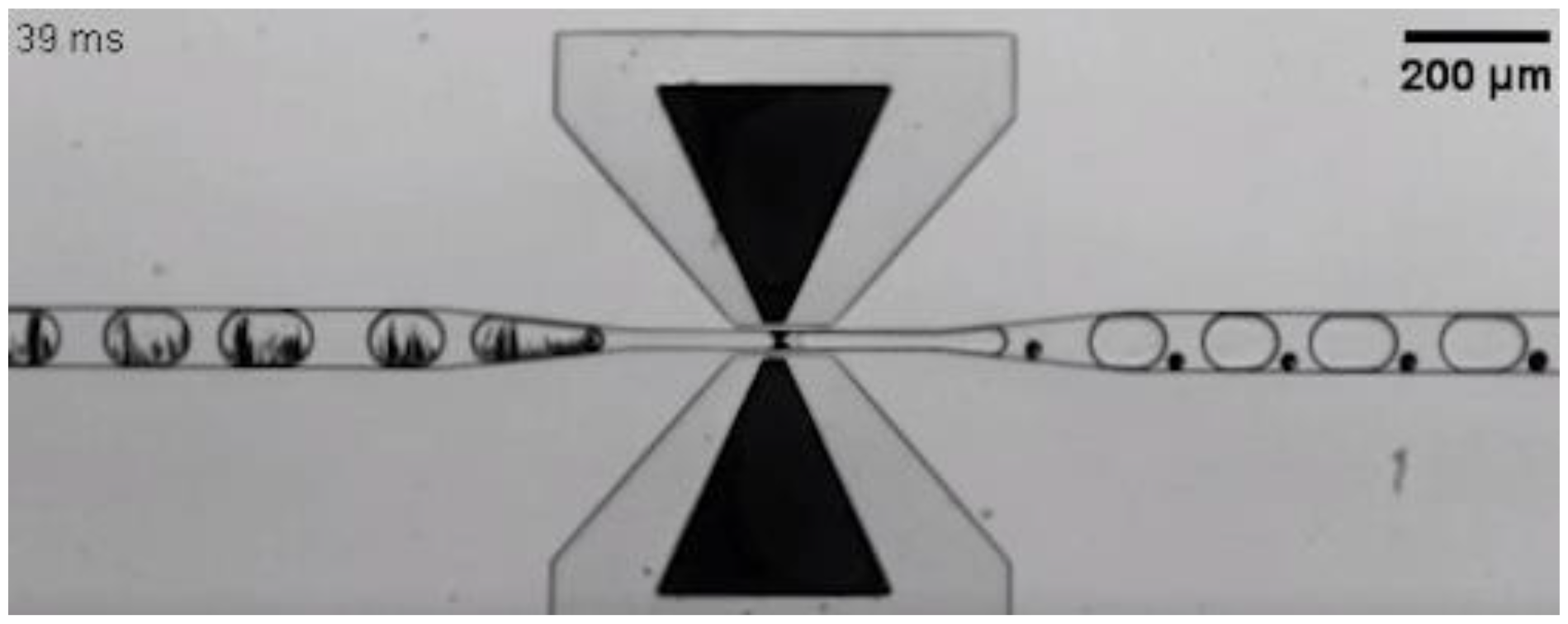
Extraction of magnetic beads from the droplets. The droplets containing beads arrive from the left. The beads are extracted as a compact cluster, which is then pushed by the following droplet.
4. When the extraction is stable, connect the outlet tubing to a 1.5 µL microcentrifuge tube filled with 300 µL carrier oil. Place the 10 mm square neodymium magnet close to the tube bottom to separate the bead cluster from the droplets (**Note 12**). Keep the collection tube on ice during extraction.

### 3.8. Assessment of mRNA purification rate by RT-qPCR

This part aims at assessing the efficiency of mRNA purification of the above protocol. The quantity of mRNA in the droplets is evaluated by RT-qPCR on the emulsion before and after extraction through the micro tweezers.

1. Using a micropipette, transfer 20 µL of the outlet emulsion in a new tube.
2. Disconnect the reinjection reservoir from the chip and dispense 20 µL of the inlet emulsion in another tube. Ensure that the two tubes contain the same volume of emulsion.
3. Add in each tube 50 µL of emulsion breaking solution. Vortex and centrifuge briefly.
4. Using a micropipette, collect 15 µL of the aqueous phase and transfer in another tube (**Note 13**). Keep the tubes on ice.
5. Prepare a serial dilution (1:10, 1:100, 1:1000) of an mRNA sample of known concentration, which will be used as a ladder to convert the qPCR Ct values to mRNA concentration.
6. Prepare the RT-qPCR master mix as follows (**Note 14**):
  ∘ 50 µL Reaction mix (2X)
  ∘ 2 µL Superscript Platinium Taq mix
  ∘ 2 µL Forward primer (10 µM)
  ∘ 2 µL Reverse primer (10 µM)
  ∘ 2 µL TaqMan probe (10 µM)
  ∘ 0.05 µL ROX dye
  ∘ 37.8 µL Nuclease free water
7. Perform RT-qPCR in reactions containing 24 µL of the above master mix with 1 µL on undiluted droplet content from step 4 (or the ladder dilution). Use the following program:
  ∘ 50 °C 15 min
  ∘ 95 °C 2 min
  ∘ 95 °C 15 s, 60 °C 30 s | x 45
8. Use a linear interpolation on the ladder Cts in log scale to get the conversion relation between the Ct and the mRNA concentration (**Figure 4A**).
9. Using this conversion, calculate the mRNA quantity in the emulsions before and after extraction to evaluate the purification efficiency (**Figure 4B**).

**Figure 4.**
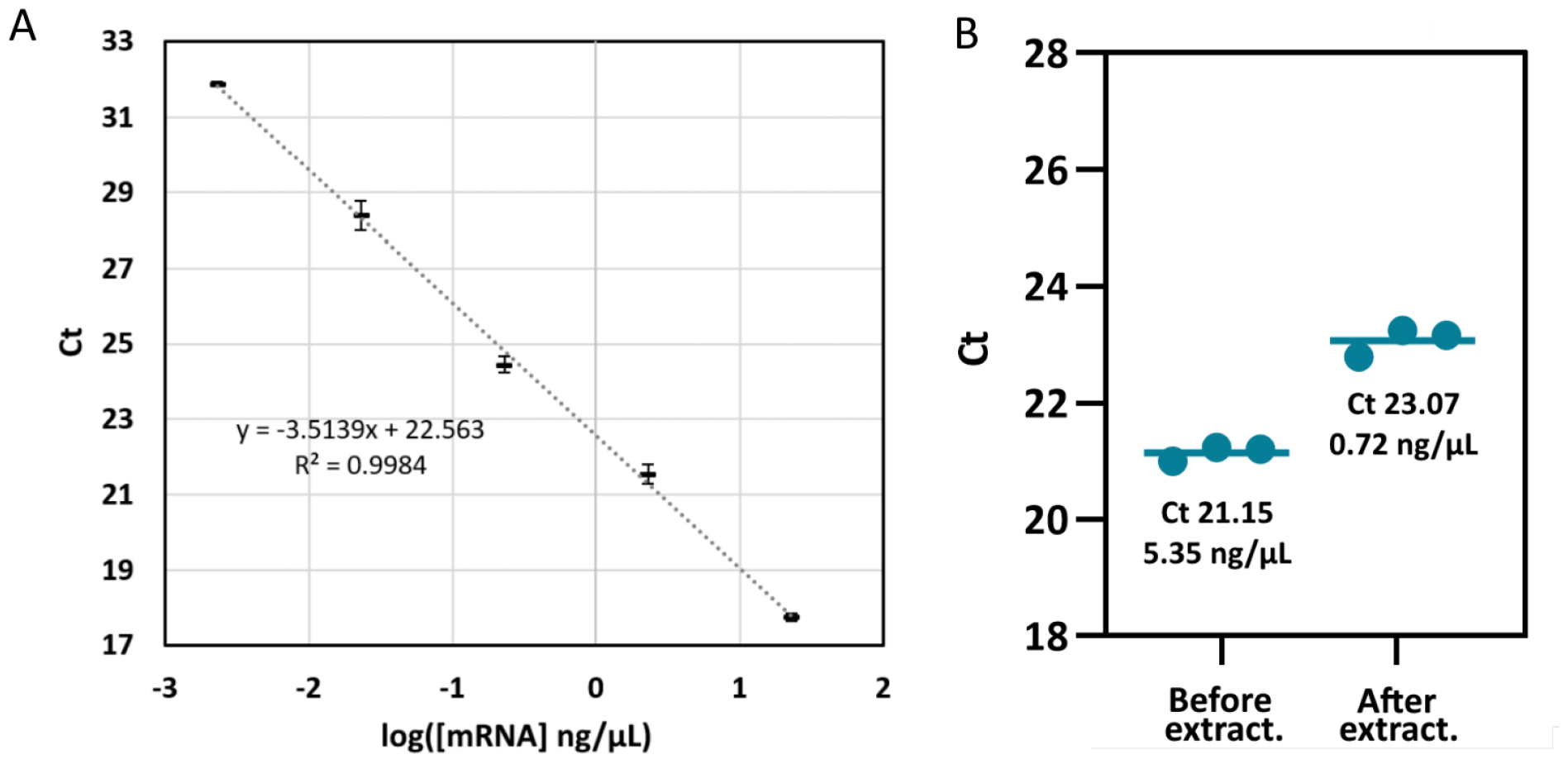
Results and analysis of RT-qPCR. (a) Calibration curve obtained with the ladder as a sample. (b) Results for the droplet sample, before and after the extraction through the micro tweezers.

## Notes

1. The deposition protocol will depend on the available equipment. We use 75 W, 15 min for Ti and 100 W, 5 min for Cu at room temperature. Some variations of seed layer thickness over experiments can be observed and are not critical. The back and edges of the substrate must be covered with tape to render only one side conductive.
2. The spin coating protocol must be optimized for each thickness. We obtained a thickness of around 50 µm with the following protocol: 300 rpm 5 s, 2600 rpm 2 s, 2800 rpm 8 s. The baking time also needs to be optimized: after cooling down, the photoresist needs to be slightly tacky but should not stick on gloves.
3. The generator must be turned on as soon as possible after immersion of the substrate in the bath to avoid copper dissolution in the acidic bath.
4. The current density is calculated with the area of patterns. The area has to be large enough (> 0.1 cm^2^) to be in the operating range of the DC generator.
5. Stirring must be strong enough to continuously renew the solution within the patterns. Use the highest stirring speed which does not induce bubbles, foaming, or instabilities.
6. The deposition may vary with the design, stirring speed, electrodes distance, bath age. A control measurement can be done after a few hours of deposition using a stylus profilometer to check the deposition rate. However, care must be taken to keep the substrate wet as much as possible as the photoresist might crack within minutes after being removed from water.
7. After a long deposition step, the water in the plating bath may have evaporated. Fill the plating bath with water back to its original level, and keep it in a bottle at room temperature.
8. The magnetic substrate can be cut into several parts if used to make several microfluidic chips. Use a laser cutter or a tungsten tip. The cut can be done before photoresist stripping to keep the substrate clean.
9. The concentrations are calculated for a final concentration in droplets of 40 µg/µL of beads and 200 cells/µL. The flow ratio between the bead and droplet solution is 2:1. This way, a cell is encapsulated for every 10 droplets of 0.5 nL.
10. When filling the syringe with the cell solution, do not use a thin needle that will cause cell lysis by shearing.
11. Beads will sediment in the tubings. The generation has to be done quickly.
12. The magnet can be locked to the tube using glue, tape, or a custom 3D printed holder. A commercial magnetic rack might be used but some are often not powerful enough.
13. In the bead containing tube, redisperse the beads homogeneously in the aqueous phase by gently pipetting up and down without disturbing the oil/water interface.
14. The master mix quantity repare as much master mix as needed, depending on the number of samples and ladder. Each reaction is preferably done in triplicates.

## 5. Conclusion

This new generation of micro-tweezers was able to create high magnetic gradients, around 20 times higher than previous technologies [14], in a microfluidic channel for the extraction of magnetic particles in sub-nanoliter droplets. Extraction of commercial 1 µm diameter magnetic particles was achieved at high throughput (20 droplets per second) with an efficiency close to 100% in 450 pL droplets. A first demonstration of its adaptability to single-cell analysis is demonstrated in this article with the extraction of mRNA (**Figure 4**). Using a purified nucleic acid solution, this unique magnetic configuration was able to reach a RNA extraction rate of 72%. This is the first demonstration of a physical separation in droplets at high throughput at single-cell scale.

## 6. Acknowledgment

The authors acknowledge all members of the IPGG technological platform for their help with microfabrication, especially Olivier Lesage for his support in photolithography and sputtering and Bertrand Cinquin for EDX spectrometry. They also thank Koceila Aizel for support with electroplating, and Fanny Tabarin and Aude Battistella for training them on cell and molecular biology (Institut Curie, UMR168), David Hrabovsky (MPBT platform, Sorbonne University) for magnetometry measurements, and Olivier Lefebvre (C2N, Univ. Paris-Saclay) for helpful advice on electroplating. This work was supported by the Institut Pierre-Gilles de Gennes (équipement d’excellence and LABEX, “Investissement d’avenir,” program ANR-10-EQPX-34). This project has received funding from the European Union’s Horizon 2020 research and innovation programme under the Marie Skodowska-Curie grant agreement No 896313.

